# Cleavage-furrow formation without F-actin in *Chlamydomonas*

**DOI:** 10.1101/789016

**Authors:** Masayuki Onishi, James G. Umen, Frederick R. Cross, John R. Pringle

## Abstract

It is widely believed that cleavage-furrow formation during cell division is driven by the contraction of a ring containing F-actin and type-II myosin. However, even in cells that have such rings, they are not always essential for furrow formation. Moreover, many taxonomically diverse eukaryotic cells divide by furrowing but have no type-II myosin, making it unlikely that an actomyosin ring drives furrowing. To explore this issue further, we have used one such organism, the green alga *Chlamydomonas reinhardtii*. We found that although F-actin is concentrated in the furrow region, none of the three myosins (of types VIII and XI) is localized there. Moreover, when F-actin was eliminated through a combination of a mutation and a drug, furrows still formed and the cells divided, although somewhat less efficiently than normal. Unexpectedly, division of the large *Chlamydomonas* chloroplast was delayed in the cells lacking F-actin; as this organelle lies directly in the path of the cleavage furrow, this delay may explain, at least in part, the delay in cell division itself. Earlier studies had shown an association of microtubules with the cleavage furrow, and we used a fluorescently tagged EB1 protein to show that at least the microtubule plus-ends are still associated with the furrows in the absence of F-actin, consistent with the possibility that the microtubules are important for furrow formation. We suggest that the actomyosin ring evolved as one way to improve the efficiency of a core process for furrow formation that was already present in ancestral eukaryotes.

## Introduction

Cytokinesis is the final stage in the cell-division cycle in which the cytoplasms and plasma membranes of the daughter cells are separated. In unikonts [animals, fungi, slime molds, and their close relatives], cytokinesis occurs by the symmetric or asymmetric ingression of a “cleavage furrow” from the periphery of the cell. For 50 years, thinking about cleavage-furrow ingression in these cells has been dominated by the contractile-actomyosin-ring (CAR) model, in which bipolar filaments of myosin-II walk along actin filaments (F-actin), much as in muscle, to produce the force that pulls the plasma membrane in to form the furrow (1–4). Actin, myosin-II, and functionally related proteins are clearly present in a ring that constricts during furrow ingression in unikont cells (5–10), and there is good evidence both that this ring produces contractile force (11) and that this force is required for normal cytokinesis in at least some cell types (12–14).

However, there are also multiple observations that are difficult to reconcile with the CAR model, at least in its simplest forms. For example, in mammalian normal rat kidney (NRK) cells, local application of the actin-depolymerizing agent cytochalasin D to the furrow region accelerated, rather than delayed, furrowing (15, 16). Moreover, equatorial furrows could form in NRK cells while myosin-II was inhibited by blebbistatin, so long as the cells were attached to a substratum (17). Additionally, motor-impaired myosin-II was able to support a normal rate of furrow ingression in mammalian COS-7 kidney-derived cells (18). In addition, myosin-II null mutants are viable and can divide some microorganisms. In the amoeba *Dictyostelium discoideum* such mutants form equatorial cleavage furrows when growing on a solid substratum (19–22), and in the budding yeast *Saccharomyces cerevisiae*, the mutant cells complete division even though they fail to assemble an actin ring at the division site (9). Although cytokinesis of the yeast mutants is inefficient, it can be almost completely rescued by expression of the myosin-II tail domain, which is incapable of generating force by myosin-actin interaction (23, 24). In a final example, modeling indicates that the actomyosin ring cannot provide more than a fraction of the force needed to drive furrow ingression in the face of intracellular turgor pressure in the fission yeast *Schizosaccharomyces pombe*, and pharmacological disassembly of F-actin after initiation of furrowing did not inhibit further furrow ingression (25).

The limitations of the CAR model become even more apparent when cytokinesis is viewed in a phylogenetic and evolutionary perspective. For example, most cells in plants divide by a mechanism (centrifugal cell-plate growth mediated by the microtubule-based phragmoplast: 26–29) that seems completely different from the cleavage-furrow ingression of unikonts, although the two groups have a common ancestor. Moreover, except for the plants, some types of algal cells, and some intracellular parasites, all non-unikont cells that have been examined divide by cleavage-furrow ingression (30–39) although they lack a myosin-II, which appears to be conserved only in the unikont lineage (40–43) [with the interesting exception of a myosin-II in the Excavate *Naegleria gruberi* (44)]. There is very little information about the mechanisms by which such furrows form, although some non-unikont cells have been reported to have actin localized in the developing furrows (33, 34, 36, 37, 45–51), raising the possibility that actin might have a role that predates and is independent of myosin-II.

Taken together, these and other observations (27, 52) suggest that the earliest eukaryotes [and the last eukaryotic common ancestor (LECA)] had a mechanism for cleavage-furrow formation that did not involve a CAR, although it might have involved actin. Importantly, such an ancestral mechanism could still exist as the underpinning for the seemingly diverse modes of cytokinesis seen today. To explore the nature of this postulated mechanism and the role of actin cytokinesis, we are studying the green alga *Chlamydomonas reinhardtii*, which divides by furrow formation (30, 53, 54; this study) but has no myosin-II (55, 56). We report here that cleavage-furrow formation in this organism does not require F-actin or any of its three non-type-II myosins. In contrast, our observations are consistent with earlier reports (30, 53, 54) suggesting a possible role for microtubules in furrow formation. Our results also suggest a previously unappreciated role for F-actin in chloroplast division.

## Results

### Live-cell observations of cleavage-furrow ingression

Previous descriptions of cytokinesis in *Chlamydomonas* have been based on light and electron micrographs of fixed cells (30, 53, 54). To observe the process in living cells, we expressed the plasma-membrane ATPase PMH1 (57) tagged with mNeonGreen (mNG) and observed cells by time-lapse microscopy. As expected, cleavage furrows ingressed primarily from one pole of each imaged cell (Fig. 1*A*, arrowhead 1; Movies S1 and S2); and reached the opposite side of the cell in 16±3 min (n=13). However, the earlier appearance of a small notch at the opposite side (Fig. 1*A*, arrowhead 2) suggested that a “lateral ingression” formed a groove around the entire perimeter of the plane of cleavage well before ingression of the medial furrow was complete, consistent with observations by DIC (Fig. 1*A*, 10′ and 12′) and electron (30) microscopy. When the imaged cells also expressed ble-GFP (a marker for the nucleus: 58), it was apparent that the medial furrow began to form at the anterior pole of the cell and progressed between the daughter nuclei (Fig. 1*B*), consistent with previous reports (30, 53). In most cells, furrow ingression was accompanied by cytoplasmic rotation (Fig. 1*A*, 0′-16′).

**Fig. 1.**
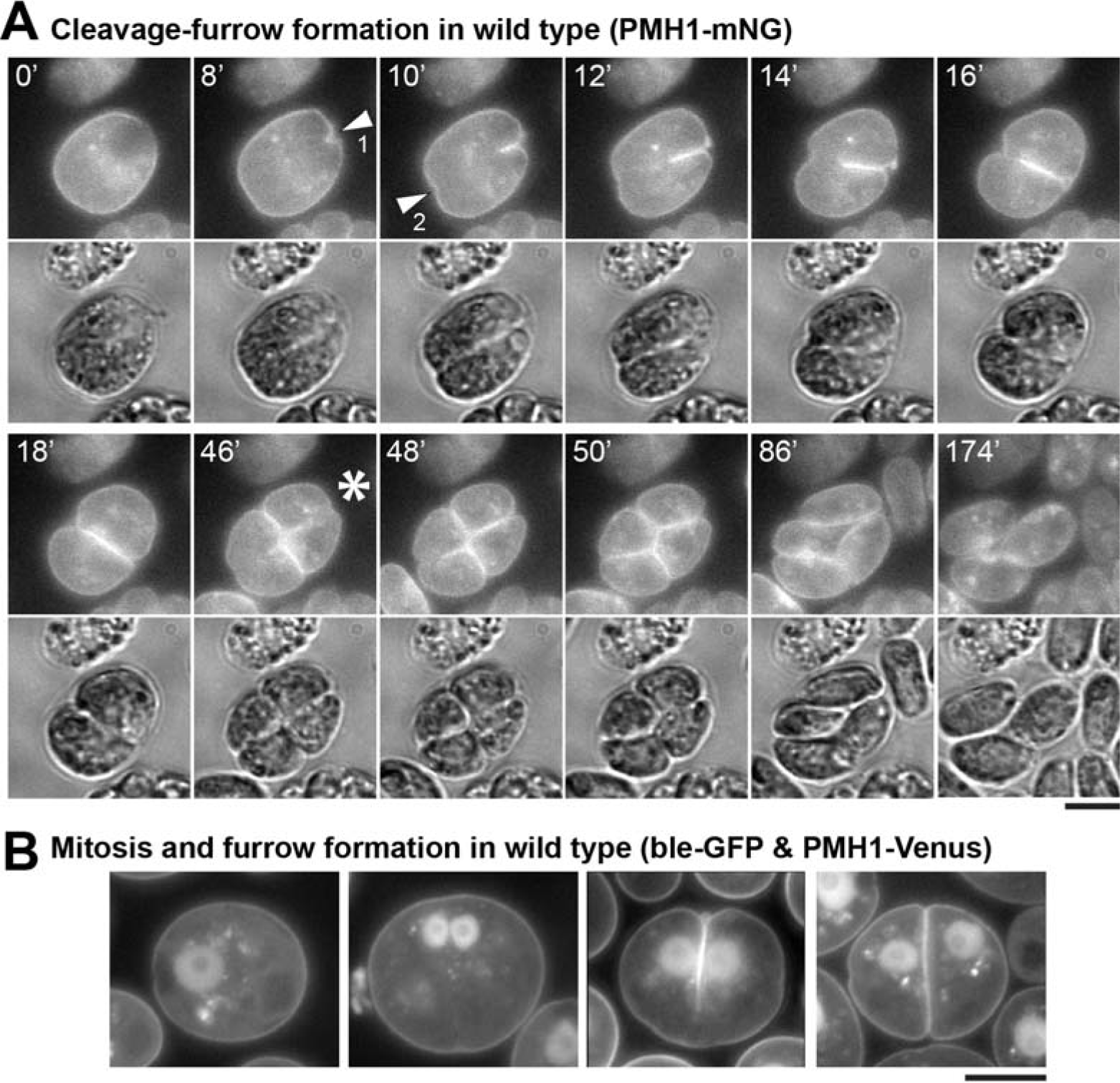
Live-cell observations of cleavage-furrow formation in *Chlamydomonas.* (*A*) Wild-type cells expressing the plasma-membrane ATPase PMH1 tagged with mNeonGreen (mNG) were synchronized using the 12L:12D/TAP agar method, mounted on TAP + 1.5% low-melting agarose, and imaged over several hours at ~25°C. Selected images are shown (times in min); the full series is presented in Movies S1 and S2. Upper images, mNG fluorescence (YFP channel); lower images, DIC. Arrowheads, positions of the initial appearance of furrow ingression visible in this focal plane in the anterior (1) and posterior (2) poles of the cell. Asterisk, onset of second cleavage. (*B*) Wild-type cells co-expressing PMH1-Venus and the nuclear marker ble-GFP were imaged using a YFP filter set during growth on TAP medium at 26°C. Cells at different stages in mitosis and cytokinesis are shown. Bars, 5 μm.

Under the growth conditions used, most cells underwent two or three rapid divisions, then hatched from the mother cell wall as four or eight daughter cells (Movie S2). The second cleavage followed the first by 38±4 min (n = 11) and was often not clearly visible because of its angle relative to the imaging plane. However, when it could be seen clearly, it always initiated at the center of the previous cleavage (Fig. 1*A*, 46′, asterisk; Movies S1 and S2).

### Localization of F-actin, but not myosins, to the cleavage furrow

Despite the lack of a type-II myosin in *Chlamydomonas*, actin could still have a role in cleavage-furrow formation (see Introduction). To explore this possibility, we expressed the F-actin-binding peptide Lifeact (59) as a fusion with mNG. In interphase cells, F-actin localized around the nucleus, at the basal body region, and in the cortex, as observed previously (Fig. 2*A*; 56, 60-62). In dividing cells, F-actin showed a transient but strong enrichment at the anterior pole (Fig. 2*B*, 0′-4′) and then appeared to be associated with the furrow throughout its ingression (Fig. 2*B*, 4′-12′; Fig. 2*C*, top). F-actin was also associated with the furrows in cells undergoing their second round of cytokinesis (Fig. 2*C*, bottom).

**Fig. 2.**
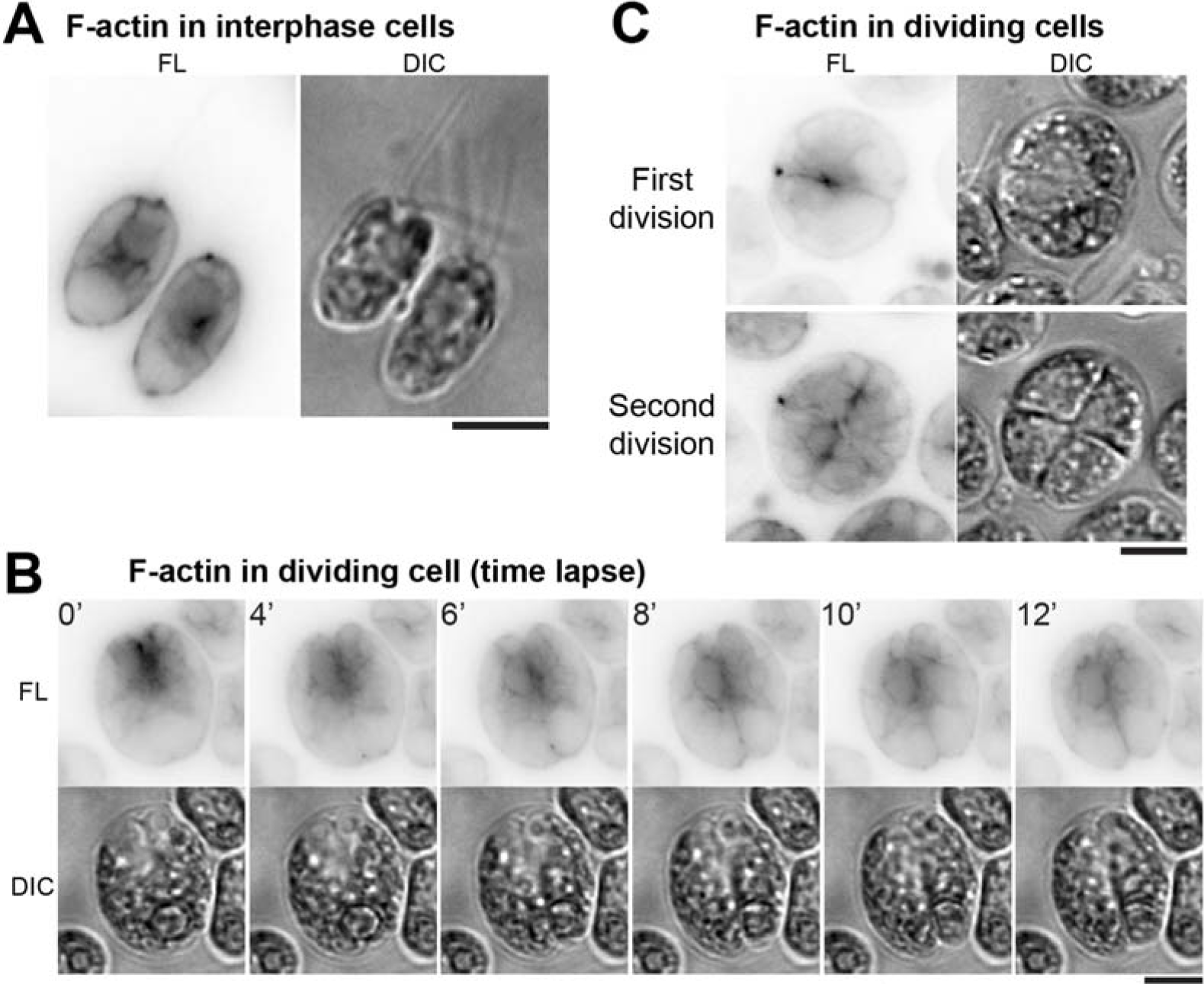
F-actin localization to the region of the cleavage-furrow. Wild-type cells expressing Lifeact-mNG were imaged; fluorescence (FL) and DIC images are shown. Fluorescence images are presented with contrast inverted for greater clarity. (*A* and *C*) Still images of interphase (*A*) and dividing (*C*) cells grown on TAP medium at 26°C. (B) Time-lapse images (as in Fig. 1*A*) showing enrichment of F-actin in the region of the cleavage furrow. Bars, 5 μm.

If the F-actin that concentrates in the furrow region has a role in cleavage-furrow formation, one or more myosins might also be involved. A BLAST search of the *Chlamydomonas* genome using the motor domain of *Drosophila melanogaster* type-II myosin detected only the three myosin genes reported previously (56). A phylogenetic analysis indicated that *MYO1* and *MYO3* encode type-XI myosins, whereas *MYO2* encodes a type-VIII myosin (Fig. 3*A*; Fig. S1*A*). Importantly, none of these myosins has an extended C-terminal coiled-coil domain such as those that allow type-II myosins to form bipolar filaments (63).

**Fig. 3.**
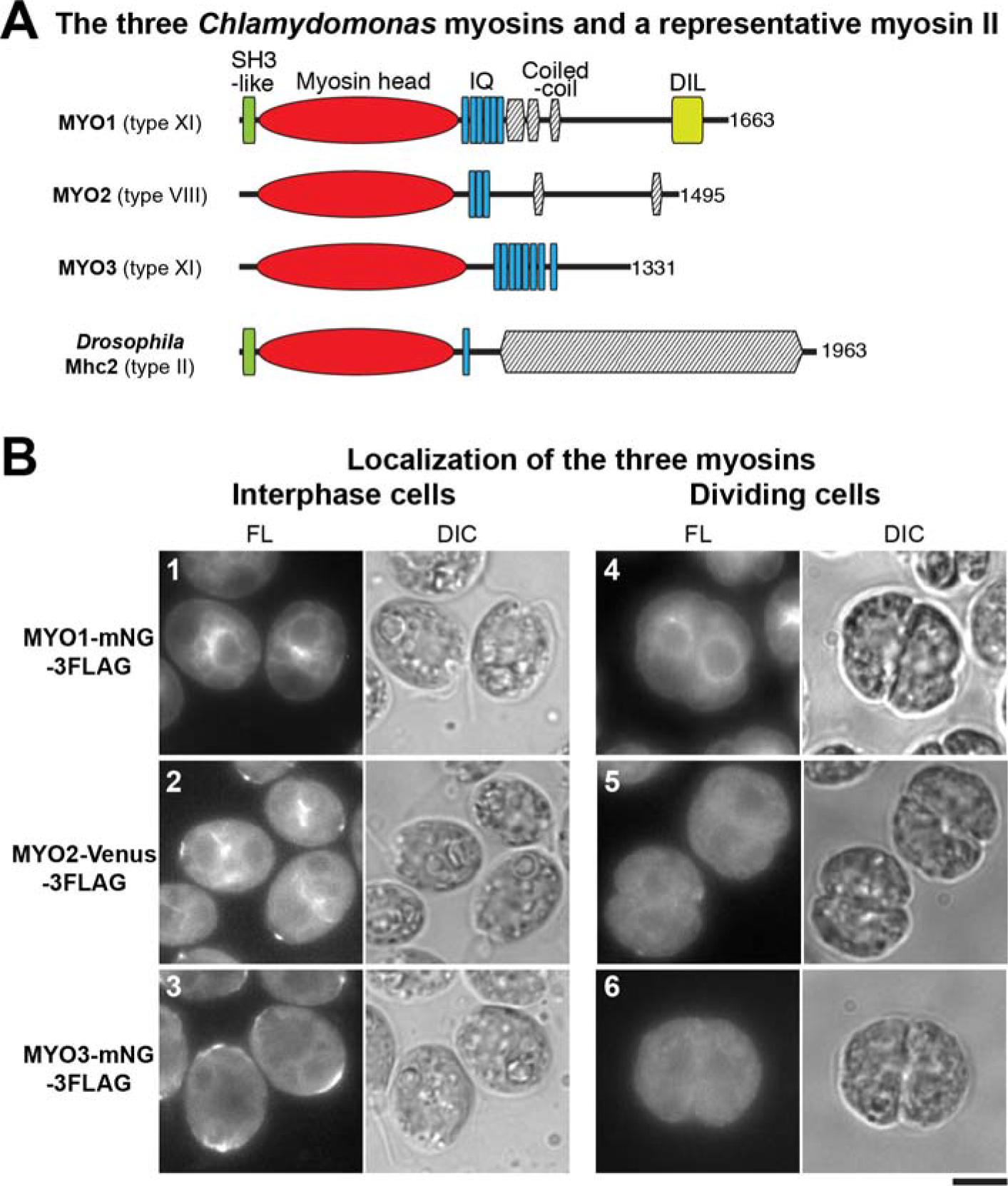
Lack of myosin localization to the region of the cleavage furrow. (*A*) Domain structures of the three *Chlamydomonas* myosins; a typical type-II myosin with a long coiled-coil tail (*Drosophila* Mhc2) is included for comparison. Domains were predicted using the HMMER (hmmer.org) and COILS (124) programs; total numbers of amino acids are indicated. (*B*) Localization of fluorescently tagged myosins in interphase and dividing cells; cells were grown on TAP medium at 26°C. Bar, 5 μm.

To ask if any of these non-type-II myosins might be involved in cytokinesis, we expressed each protein in wild-type cells with mNG-3FLAG or Venus-3FLAG fused at its C-terminus; in each case, the fusion protein was detected at or near the expected molecular weight by Western blotting (Fig. S1*B*). In interphase cells, MYO1 was enriched in the perinuclear region (Fig. 3*B*, 1; Fig. S1*C*, 4) in a pattern overlapping one subdomain of F-actin localization as seen with the tagged Lifeact probe (Fig. 2*A*; Fig. S1*C*, 1). MYO2 localized to the perinuclear region as well as to dots at the cell-anterior region near the basal bodies (Fig. 3*B*, 2; Fig. S1*C*, 7), again resembling aspects of F-actin localization. MYO3 localized to the cell-anterior region, as well as to the cortex in the cell-posterior (Fig. 3*B*, 3; Fig. S1*C*, 10). Importantly, in dividing cells, none of the myosins showed any detectable enrichment around the cleavage furrow (Fig. 3*B*, 4-6).

The similarity in localization of the myosins to that of F-actin in interphase cells (see above) suggests that the tagged myosins interact normally with actin. Further evidence for this conclusion was obtained in experiments that exploited the *Chlamydomonas* system for F-actin homeostasis (60, 61, 64, 65). In vegetative wild-type cells, only the conventional actin IDA5 is expressed. Exposure of cells to the F-actin-depolymerizing drug latrunculin B (LatB) leads to a rapid disassembly of F-IDA5 (Fig. S1C, 2), degradation of the monomeric IDA5, and upregulation of the divergent actin NAP1, which provides actin function by assembling into LatB-resistant filaments (Fig. S1C, 3). Similarly, the localization signals for MYO1 and MYO3 were largely lost during a short incubation with LatB but subsequently recovered (Fig. S1C, 4-6 and 10-12), suggesting that these tagged myosins can bind to both F-IDA5 and F-NAP1. The perinuclear signal for MYO2 was also sensitive to LatB but did not recover (Fig. S1C, 7-9; 56), suggesting that this myosin binds only to F-IDA5.

Thus, although the current lack of null mutations for any of the myosin genes precludes a definitive test of the function of the tagged proteins, it seems most likely that they at least localize as expected in a F-actin or NAP dependent manner, so that their apparent absence from the furrow region suggests that any function of actin in furrowing does not involve the myosins (e.g., in a noncanonical actomyosin ring).

### Cleavage-furrow ingression in the absence of F-actin

To ask whether actin (with or without myosin) plays a role in cleavage-furrow formation, we took advantage of our prior isolation of a null mutation (*nap1-1*) in the *NAP1* gene (60). Because F-IDA5 is highly sensitive to LatB, treatment of a *nap1-1* strain with the drug results in a rapid and complete loss of F-actin as detected by LifeAct. LatB treatment is lethal for *nap1-1* mutants, indicating that F-actin is directly or indirectly important for various cellular processes in *Chlamydomonas*. We found previously that LatB treatment of *nap1-1* cells blocked cell growth, and DNA replication and cell division were also blocked, likely as an indirect consequence of the block to cell growth. To evaluate a possible specific role of actin in cytokinesis, we employed an accurate cell-cycle synchronization method using a 12:12 light-dark cycle (66, 67). Under the conditions used, the cells grew in size throughout the light phase, began to divide at ~13 h (i.e., 1 h into the dark phase), and hatched out as small daughter cells at ≤20 h (Fig. 4*A*, top row). We then treated aliquots of the culture with LatB at different times before and during the onset of cytokinesis and examined the cells several hours later (Fig. 4*A*, bottom row). When LatB was added at ≤9 h, the cells ceased growth and never detectably initiated cytokinesis. Flow-cytometry analysis in separate but similar experiments indicated that most of these cells had arrested before DNA replication as expected (60). When LatB was added at 10 h, most cells remained round and did not begin furrow formation, but a few formed what appeared to be normal cleavage furrows (Fig. 4*A*, arrowhead) or “notch”-like structures (Fig. 4*A*, arrow). A similar experiment using cells expressing PMH1-Venus and ble-GFP indicated that the cells with notches had not undergone mitosis (Fig. S2*A*, left). In contrast, when LatB was added at ≥11 h, many cells appeared to have gone through two or more rounds of cleavage-furrow ingression, forming clusters of 4-8 cells, each of which contained a nucleus (Fig. 4*A*; Fig. S2*A*, right). In a separate but similar experiment, we noted a rough positive correlation between size of the undivided cells and their likelihood of forming a furrow in the presence of LatB (Fig. S2*B*), suggesting either a direct size requirement for cytokinesis or a requirement for some size-correlated cell-cycle event for actin-independent furrow formation.

**Fig. 4.**
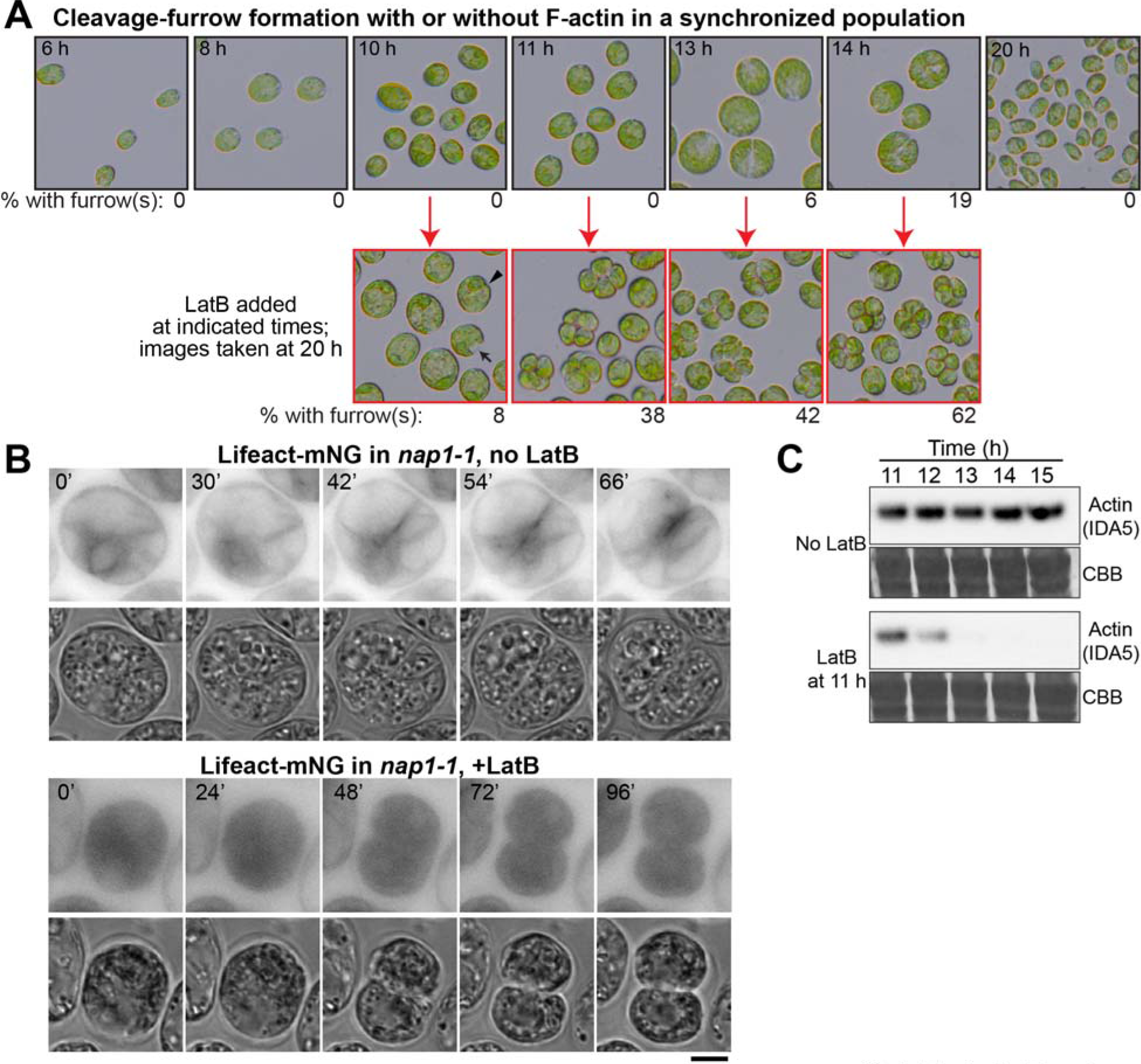
Cleavage-furrow formation in the absence of F-actin. (*A*) *nap1-1* cells were synchronized using the 12L:12D/liquid TP method at 26°C, incubated for up to 20 h, and imaged at intervals (top row). At the indicated times (red arrows), samples were plated on TAP agar containing 3 μM LatB, incubated at 26°C, and then imaged at 20 h (i.e., 8 h into the dark period). The percentages of cells with visible cleavage furrows are shown below the images. (*B*) F-actin localization in *nap1-1* cells with or without LatB treatment. *nap1-1* cells expressing Lifeact-mNG were synchronized using the 12L:12L/TAP agar method at 26°C, mounted on TAP + 1.5% low-melting agarose with or without 3 μM LatB, and observed during growth at 26°C. Selected images are shown; contrast is inverted for greater clarity. The LatB-treated cell had been incubated with the drug for ~4.5 h before the first frame shown. Note that dispersion of Lifeact-mNG into the entire cytoplasm contributed to the strong apparent background in these cells. Bar, 5 μm. (*C*) Rapid degradation of IDA5 upon LatB addition to a synchronized culture. *nap1-1* cells were synchronized as in (*A*) and the culture was split. 3 μM LatB was added to one culture at 11 h, and samples were drawn at the indicated times and subjected to Western blotting using an anti-actin antibody. 30 μg total protein were loaded in each lane. CBB, the membrane stained with Coomassie Brilliant Blue, shown as a loading control.

Although our prior work had suggested that LatB-treated *nap1-1* cells contained no residual F-actin, it seemed possible that there might be a special population of drug-resistant filaments in the cleavage-furrow region. However, no such filaments were observed when time-lapse observations were made on LatB-treated *nap1-1* cells expressing Lifeact-mNG (Fig. 4*B*). Moreover, consistent with our prior observations indicating rapid proteasomal degradation of LatB-depolymerized IDA5 (61), IDA5 was also largely or entirely degraded by the time such cells began furrow formation (Fig. 4*C*).

### Reduced efficiency of furrow ingression in cells lacking F-actin

To investigate the efficiency of cleavage-furrow formation in the absence of F-actin, we performed time-lapse microscopy on *nap1-1* cells expressing PMH1-mNG. In the absence of LatB, furrow formation in these cells proceeded to completion in 17±4 min (n=12) (Fig. 5, *A* and *C1*; Fig. S2*C*, 2), not significantly different from the rate in wild-type cells (16±3 min; n=13) (Fig. 1; Fig. 5*C1*; Fig. S2*C*, 1). In contrast, in the presence of LatB, although the rates of furrow ingression varied considerably in individual cells, they were slower in all cells examined than in control cells even during the early stages of furrow ingression, and more so during its later stages (Fig. 5*B*, *C1*, *C2*; Fig. S2*C*, 3). The time gap between the formation of a small cell-anterior notch and the detectable ingression of the medial furrow was expanded from ~2 min in control cells to 5-15 min. Moreover, although the medial furrow typically ingressed smoothly into about the middle of the cell, the second notch at the cell-posterior end did not appear normally at that time, and, in most cells, the medial furrow stalled at that point for an extended period before eventually appearing to complete its growth (10 of the 12 cells examined: Fig. 5*B1*, arrow; Movie S4) or regressing (one of the 12 cells examined: Fig. 5*B2*; Movie S5). More rarely (one of the 12 cells examined), the furrow appeared to progress across the cell without interruption (Fig. 5*B3*; Movie S6). In all cells examined, the daughter cells remained clustered without hatching, and the fluorescence of PMH1-mNG became quite dim, making it difficult to determine when (or whether) the plasma membranes were fully resolved. Indeed, when the clusters of LatB-treated *nap1-1* cells (see Fig. 4*A*) were treated with the cell-wall digesting enzyme autolysin, ~5% of the cells remained connected with an intercellular bridge between the pairs (Fig. S2*D*). Taken together, these results suggest that although actin is not required for furrow ingression per se, it plays some ancillary role(s) that facilitates the early stages of furrowing and become(s) more important during the later stages of furrow ingression and/or abscission.

**Fig. 5.**
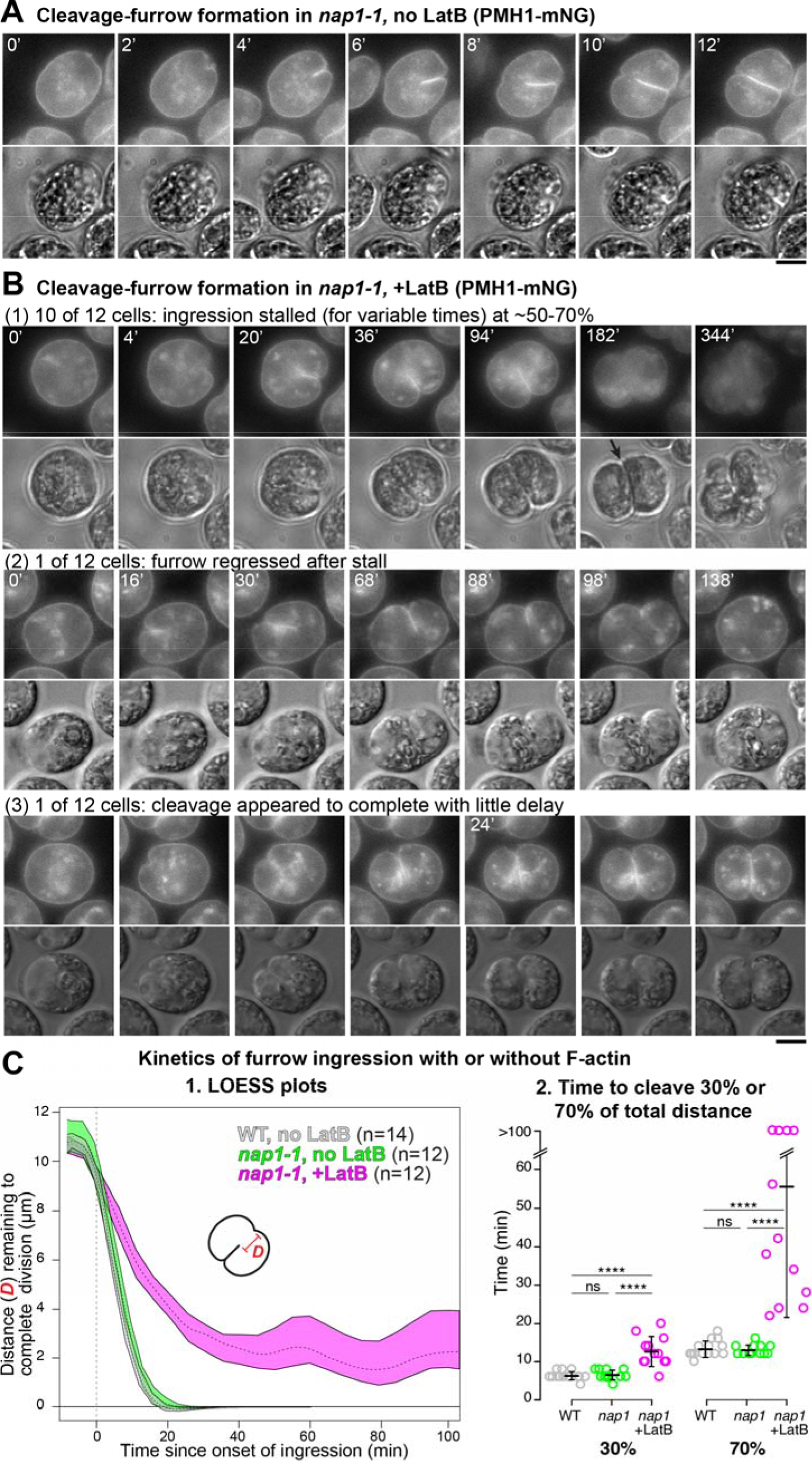
Slower cleavage-furrow ingression and delay in furrow completion in the absence of F-actin. (*A*) *nap1-1* cells expressing PMH1-mNG were synchronized using the 12L:12D/TAP agar method and observed by time-lapse microscopy at 26°C. Full series is presented in Movie S3. Bar, 5 μm. (*B*) As in *A* except that 3 μM LatB was added 120-160 min before the first frames shown. Selected images are shown to illustrate the different time scales; full series are presented in Movies S4-S6. Bar, 5 μm. (*C*) Kinetics of furrow ingression in wild-type cells without LatB (Fig. 1*A*, Movie S2) and in *nap1-1* cells with or without 3 μM LatB. Data are from the experiments shown in *A* and *B*. (1) Distance of the leading edge of the medial furrow from the opposite side of the cell as a function of time since the onset of furrow ingression (set at 0). Means and 95% confidence intervals of 1000X-bootstrapped LOESS curves are shown (see Materials and Methods). The curves for individual cells are shown in Fig. S2*C*. (2) Times for the furrow to reach 30% or 70% of the total distance across the cells. Because of mNG bleaching after prolonged time-lapse observations, cleavage times were capped at 100 min for these analyses. Bars indicate means and standard deviations. Statistical analyses were performed using one-way ANOVA and Tukey’s post-hoc multiple comparisons (ns, not significant; ****, *P*<0.0001).

### Association of microtubules with the cleavage furrow in the absence of F-actin

It has long been known that microtubules (MTs) are associated with the cleavage furrows in *Chlamydomonas*, and their depolymerization by drugs or mutation largely blocks cytokinesis (30, 53, 54). To ask if this association is maintained in the absence of F-actin, we first tried, but failed, to visualize the furrow-associated MTs by time-lapse imaging using a fluorescently tagged tubulin. However, we had better results upon expressing an mNG-tagged version of the plus-end-binding protein EB1 (68). In both wild-type cells (68) and *nap1-1* cells not treated with LatB (Fig. 6*A*, 0′; Movie S7), EB1-mNG was concentrated in the basal-body region of pre-mitotic cells. Upon entry into mitosis, EB1-mNG disappeared from the cell pole and appeared in the mitotic spindle (Fig. 6*A*, 4′-6′; Movie S7). After mitosis, the EB1-mNG signal reappeared at the apical pole of the cell, from which it moved into and across the cell body as the cleavage furrow formed (Fig. 6*A*, 12′-14′; Movie S7) and remained concentrated in the middle of the division plane after cytokinesis (Fig. 6*A*, 20′; Movie S7). Each daughter cell then formed an EB1-labeled spindle in the region proximal to this site, and the new furrows (as marked by EB1-mNG) grew outward from the center of the cell to the surface (Fig. 6*A*, 44′ and 48′, arrows). In *nap1-1* cells treated with LatB, EB1-mNG localization was nearly normal during the first division (Fig. 6*B*, 0′-22′; Movie S8), except for a slower-than-normal progression through the cytoplasm, consistent with the PMH1-mNG data. However, a fraction of the EB1-mNG appeared to leave the division plane prematurely and form foci at the far sides of the daughter cells (Fig. 6*B*, 58′-126′, arrowheads), suggesting a defect in polarity maintenance caused by the loss of F-actin. Probably as a consequence, the spindles for the second mitosis were positioned distal from the previous division plane, and the cleavage furrows subsequently grew inward (Fig. 6*B*, 152′-164′, arrows; Movie S8), although the localization of EB1-mNG, and thus presumably of both the spindle and furrow-associated MTs, otherwise appeared essentially normal. This preservation of nearly normal association of MTs with the furrows is consistent with the hypothesis that the MTs may be involved in furrow ingression both in the presence and absence of F-actin.

**Fig. 6.**
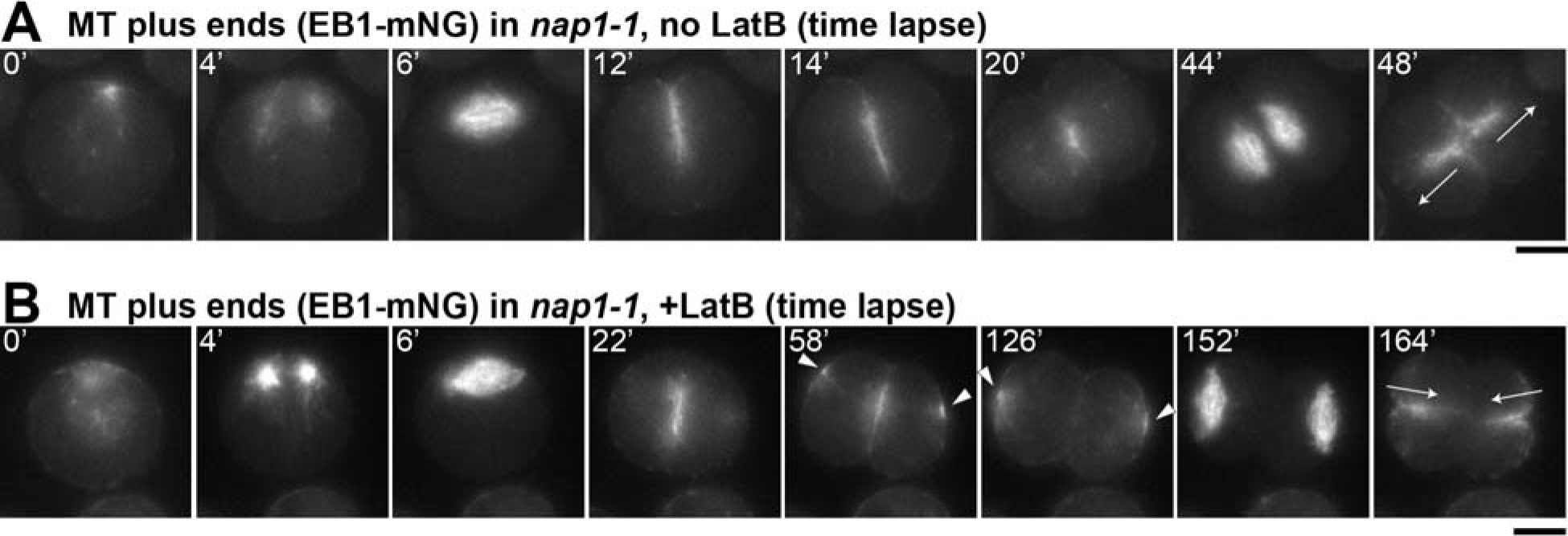
Persistent association of microtubules with cleavage furrows, but changes in polarity during the second division, in the absence of F-actin. *nap1-1* cells expressing EB1-mNG to visualize microtubule plus ends were synchronized using the 12L:12D/TAP agar method and observed by time-lapse microscopy at 26°C. Selected images are shown; full series are presented in Movies S7 and S8. (*A*) A cell not treated with LatB. (*B*) A cell treated with 3 μM LatB beginning ~20 min before the first frame shown. Arrowheads, aberrant foci of EB1-mNG at positions distal to the cleavage furrow in cells lacking F-actin; arrows, the directions of furrowing during the second division of each cell. Bars, 5 μm.

### Defective chloroplast division in cells without F-actin

The EB1-mNG studies also revealed a possible cause of the delay in furrow completion in cells lacking F-actin. In normal *Chlamydomonas* cells, the large, cup-shaped chloroplast is centered on the posterior pole of the cell (Fig. 7*A*, 0′), so that the organelle lies squarely in the path of the ingressing cleavage furrow. In control cells, division of the chloroplast (as visualized by chlorophyll autofluorescence) appeared to have occurred by the time that the furrow (or at least its associated EB1) reached the organelle (Fig. 7*A*, 3′, arrowhead), confirming previous observations that the chloroplast has a division machinery that is independent of, although temporally and spatially coordinated with, cleavage-furrow ingression (69, 70). In contrast, in cells lacking F-actin, the furrow appeared to partially penetrate into the undivided chloroplast over an extended period (Fig. 7*B*, 21′-111′), before finally moving through a gap formed in the chloroplast (Fig. 7*B*, arrowhead). These results suggest that efficient chloroplast division requires F-actin, that coordination of division between the cell and the plastid requires F-actin, or both, and that the physical barrier posed by the undivided chloroplast may explain the delay in furrow completion in cells lacking F-actin.

**Fig. 7.**
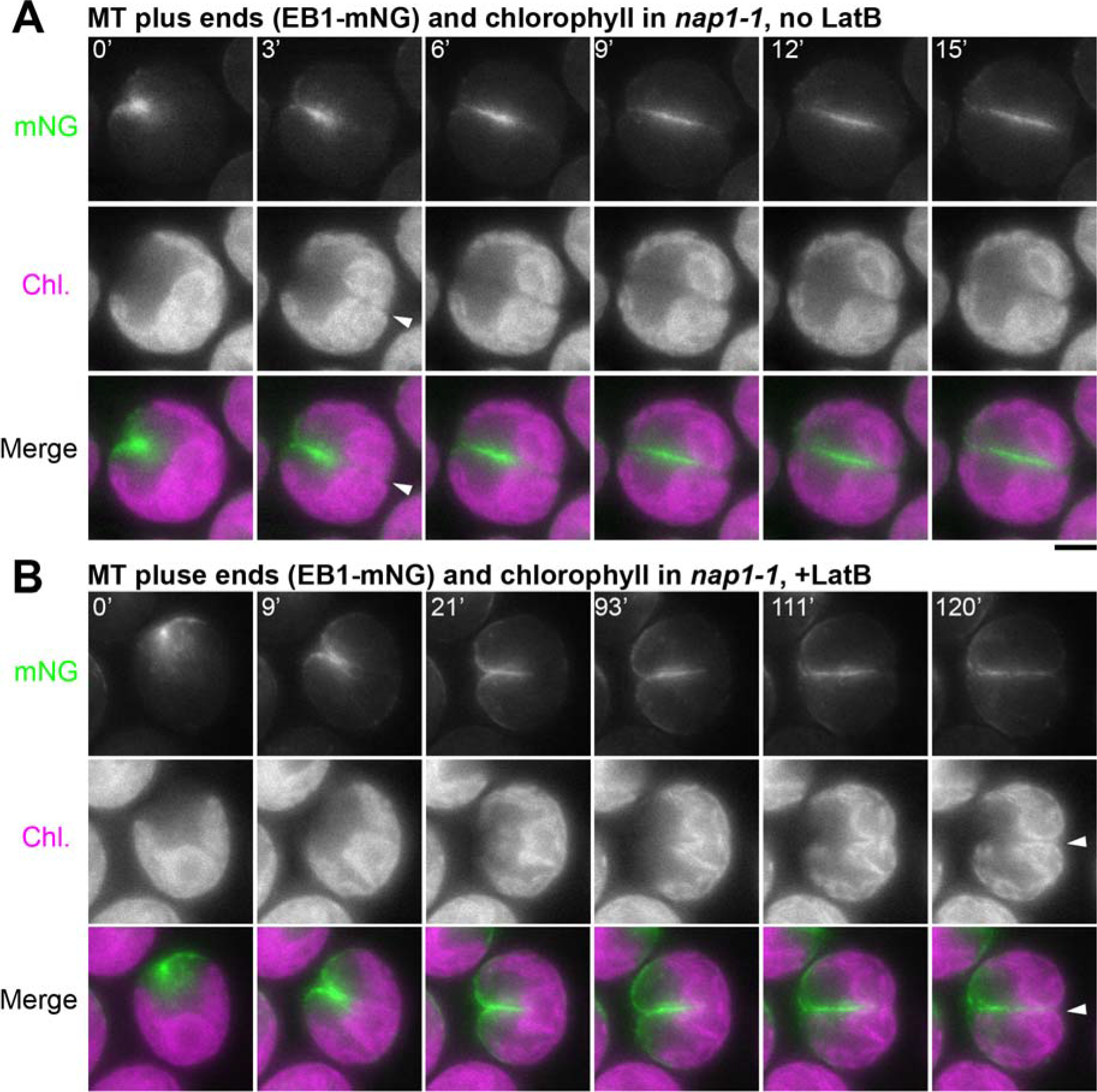
Delayed chloroplast division in the absence of F-actin. EB1-mNG fluorescence and chlorophyll autofluorescence are shown in a cell not treated with LatB (*A*) and a cell treated with 3 μM LatB beginning ~20 min before the first frame shown (*B*). Cells had been synchronized prior to imaging using the 12L:12D/TAP agar method. Arrowhead, time of apparent completion of chloroplast division. Bars, 5 μm.

## Discussion

### Rate of cleavage-furrow ingression

Both electron microscopy (30, 53, 71) and light microscopy (54, 72) had suggested that *Chlamydomonas* divides by means of an asymmetrically ingressing cleavage furrow (67), despite its lack of a type-II myosin, and we confirmed this model by live-cell imaging using a fluorescent plasma-membrane marker. The time-lapse imaging also allowed us to determine the rate of furrow ingression. At its maximum (~0.85 μm/min; see Fig. 5*C*), this rate is comparable to those in small- to medium-sized cells that have type-II myosins and form actomyosin rings at their furrow sites, such as *S. pombe* (~0.15 μm/min: 25, 73, 74), *Neurospora crassa* (1.3-3.2 μm/min: 75), and various mammalian somatic cells (3-4 μm/min: 18, 76-79). Thus, the presence of a CAR is not necessary to produce a rate of furrow ingression in this range.

### Localization of F-actin, but not myosin, to the cleavage furrow

Our live-cell imaging also clarified the spatial relationships of the furrow, F-actin, and the three *Chlamydomonas* myosins. Although immunostaining had indicated that there was actin in the furrow region (47, 54, 65, 80), it was not clear whether this actin was in filamentous form (47). However, imaging of cells expressing the F-actin-specific probe Lifeact showed clearly that F-actin is enriched in the furrow region during most or all of the period of furrow ingression. In contrast, although a previous report had suggested (based on immunostaining with an antibody to *Dictyostelium* type-II myosin) that myosin is also localized to the furrow region (54), fluorescence tagging of the three *Chlamydomonas* myosins (two type XI and one type VIII) showed no enrichment in this region. Although this conclusion should be qualified by the lack of definitive evidence that the tagged myosins were fully functional, the loss of their localization after LatB treatment suggests that they co-localize normally with F-actin (see Results). Moreover, the likelihood of a myosin role in *Chlamydomonas* furrow formation is also reduced by our finding that F-actin itself is not essential for this process.

### Cleavage-furrow ingression without F-actin

Despite the apparent absence of myosins from the furrow region in *Chlamydomonas*, it seemed possible that the F-actin there might play an essential role in furrow formation. However, when we used a combination of a mutation (to eliminate the drug-resistant actin NAP1) and a drug (to depolymerize the drug-sensitive actin IDA5), the cells were still able to form cleavage furrows and divide. While it is theoretically possible that a population of drug-resistant IDA5 filaments remained specifically in the cleavage furrow, we could detect no such filaments by Lifeact staining. Moreover, Western blotting revealed that, as observed previously in asynchronous cells (61), depolymerization of F-IDA5 in dividing cells was followed quickly by degradation of IDA5 itself. Taken together, our results appear to establish that the actin cytoskeleton does not play a principal role in generating the force for cleavage-furrow ingression in *Chlamydomonas*.

Nonetheless, several observations indicate that actin does contribute significantly to cell division in *Chlamydomonas*. First, in the absence of F-actin, the rate of early furrow ingression was ~2-fold slower than normal. It seems possible either that actin forms a contractile structure, not dependent on myosin, that contributes some force for furrow ingression or that actin is necessary for the trafficking of Golgi-derived vesicles that ultimately provide the new cell surface material for the growing furrow. Second, the last ~30% of cleavage was slowed even more (≥5 fold), and some cells appeared to fail (or at least have long delays in) the final abscission of the daughters. In addition, EB1 signal (marking MT plus-ends) disappeared prematurely from the furrow region when F-actin was missing, presumably reflecting a failure to maintain MT organization during the final phase of cleavage. Finally, the directions of the second cleavages were inverted, probably because the polarities of the two daughter cells produced by the first cleavage were also inverted. The precise mechanisms by which F-actin facilitates cytokinesis are not yet clear, but the observed slowness of late furrow ingression in its absence appears to at least partly reflect a delay in chloroplast division (see below). Together, our findings highlight the value of using *Chlamydomonas* to explore non-CAR-related roles of the actin cytoskeleton in cytokinesis.

### An apparent role for F-actin in chloroplast division

In most cells lacking F-actin, there was a delay in chloroplast division of ≥120 min. This observation was surprising, because despite early reports of an actin role in plastid division in both charophyte and red algae (81, 82), no such role has been recognized (to our knowledge) in plants or other organisms. As the *Chlamydomonas* chloroplast lies directly in the path of the ingressing cleavage furrow, the delay in chloroplast division may explain much or all of the delay also seen in furrow completion in these cells. Although we cannot currently test this model due to the lack of a method for clearing the chloroplast from the division path, a similar apparent obstruction of cytokinesis by an undivided chloroplast was observed after expression of a dominant-negative dynamin mutant in the red microalga *Cyanidioschyzon merolae* (83). Most other unicellular algae also contain only one or a few chloroplasts, whose division must presumably be coordinated both temporally and spatially (i.e., division in the same plane) with that of the cell (69). The temporal coordination is achieved, at least in part, by cell-cycle control of the expression of the proteins (such as FtsZ and dynamin) directly involved in chloroplast division (66, 84). However, the spatial coordination seems to require a local, structure-based signal. Our results suggest that the furrow-associated F-actin may provide this spatial cue to the chloroplast-division machinery and, in so doing, might also dictate the precise timing of its action.

### Possible role for MTs in cleavage-furrow formation

It has long been thought that MTs may be involved in cleavage-furrow positioning and/or ingression in *Chlamydomonas*. Electron-microscopy and immunofluorescence studies have shown that two of the four “rootlet” MTs that run from the basal body along the cortex align with the division plane (like the preprophase band in plant cells), while an array of many MTs, the “phycoplast”, runs along the developing furrow (30, 53, 54, 85, 86). Moreover, pharmacological disruption of the MTs inhibits cytokinesis (54). Our observations on cells expressing a fluorescence-tagged EB1 protein have now added the information that dynamic MT plus-ends are associated with the furrow throughout its ingression and that this association is maintained even during furrow formation in the absence of F-actin. Thus, it seems likely that the MTs have a direct, actin-independent role in promoting furrow ingression, possibly by guiding the deposition of new cell-surface materials in the growing furrow as they do in plant-cell phragmoplasts. Testing this hypothesis and elucidating the mechanisms involved will be major goals of future studies.

### Cytokinesis in phylogenetic and evolutionary perspective

Our study also sheds some light on the evolutionary origins and underlying basal mechanisms of eukaryotic cytokinesis. The wide phylogenetic distribution of division by cleavage-furrow ingression, together with the near-universal absence of myosin-II outside the unikonts, had already made clear that a conventional CAR model cannot account generally for furrow formation. Moreover, we have shown here that even F-actin is not essential for the formation of cleavage furrows in *Chlamydomonas*. This observation has some precedents and parallels. Even within the unikonts, it is clear that F-actin is not essential for furrow ingression in many cases (9, 15, 25, 87, 88; and see Introduction), and this may be the rule, rather than the exception, in non-unikonts. For example, in the ciliate *Tetrahymena pyriformis*, latrunculin A did not block cell division despite a loss of F-actin and a consequent disruption of actin-dependent processes such as food-vacuole formation (39); in the Diplomonad parasite *Giardia lamblia*, which has one actin but no myosin(38), furrowing occurred efficiently when actin expression was knocked down with a morpholino (89), although the cells were delayed in abscission; and in the red alga *C. merolae*, which also has no myosin, cells divide by furrowing even though actin is not expressed under normal growth conditions (90). Taken together, these and related observations suggest that in the earliest eukaryotes, the LECA, and most branches of the modern eukaryotic phylogeny, actin and myosin are not primarily responsible for the force that produces cleavage-furrow ingression. We speculate that as prominent roles for actin and myosin evolved in the unikonts, the underlying ancestral mechanisms for driving furrow ingression may remain in force.

In thinking about these ancestral mechanisms, it is instructive to consider the mechanisms of cytokinesis in modern prokaryotes. Bacteria divide using a furrowing mechanism in which the tubulin-like FtsZ plays a central role (91–95), which is likely to be used also by many archaea (96–100), so the immediate prokaryotic ancestor of the first eukaryotes probably also divided by such a mechanism. Current information about FtsZ action suggests that it functions both to bend the inner membrane and to organize the symmetric deposition of cell wall, which drives in the membrane to produce the division furrow (93–95, 101). Thus, if an ancient FtsZ was the evolutionary progenitor of modern tubulin, the major role of MTs in cleavage-furrow formation in such distantly related modern eukaryotes such as *Chlamydomonas, G. lamblia* (89), *Penium margaritaceum* (102)*, Trypanosoma brucei* (103)*, Tetrahymena thermophila* (104), and *Toxoplasma gondii* (105) may reflect the persistence of an ancient mechanism for adding cell-surface material to form a furrow. The same interpretation might apply even to the association of parallel arrays of MTs with cleavage furrows observed in some animal cells, such as embryos of *Xenopus* (106, 107), zebrafish (108), and *Drosophila* (109), or the midbody MTs formed during metazoan abscission (27), where the MTs appear to play an important role in targeting vesicles containing new membrane. In any case, the hypothesis of a primordial role for MTs in eukaryotic cytokinesis seems to make it easier to understand the central role of MTs in cytokinesis in modern plants, where both the preprophase band (which marks the future division plane) and the phragmoplast (which organizes the centrifugal deposition of new cell membrane and cell wall by fusion of post-Golgi vesicles) are MT based (29).

At the same time, it should also be noted that there is now good evidence that the prokaryotic ancestor of modern eukaryotes also had an actin-like protein (110, 111), and there is even some evidence that this protein might have been associated with division sites (112) despite the lack of evidence for any myosin in such organisms. Thus, the association of actin with the furrow regions both in *Chlamydomonas* and in many other eukaryotes without a myosin II may also be a preserved ancestral trait. In this regard, it is interesting that the preprophase band in plants involves actin as well as MTs, conceivably reflecting an earlier stage in plant evolution in which MTs and actin functioned together to bring about ingression of a furrow (113). If both the division mechanisms of modern plants and those of modern unikonts evolved from such an ancestral state (by recruitment of intracellular MTs to form the phragmoplast, and by reduction of the MT role in furrow formation in favor of an actomyosin system, respectively), then continuing studies of *Chlamydomonas* should help to elucidate both the two evolutionary paths and both of the modern mechanisms.

In summary, we suggest that a full understanding of eukaryotic cytokinesis, even in the intensively studied animal cells, will remain elusive unless a greater effort is made to incorporate the lessons about the evolution of this process that can be learned by studying it in the full diversity of modern eukaryotes.

## Materials and Methods

### Strains, growth conditions, and genetic analysis

*C. reinhardtii* wild-type strains CC-124 (mt-) and iso10 (mt+, congenic to CC-124) were the parental strains. The *nap1-1* mutant had previously been isolated and backcrossed three times in the CC-124 background (60). The ble-GFP and PMH1-Venus strains are progeny of previously established transgenic strains (57, 58, 114).

Routine cell culture was done in Tris-acetate-phosphate (TAP) medium (115) at ~26°C under constant illumination at 50-100 μmol photons m^−2^ s^−1^. The same medium without acetate (TP) was used in one method for cell-cycle synchronization (see below). Except for synchronized cultures, liquid cultures were in exponential phase when experiments were performed. LatB was purchased from Adipogen (AG-CN2-0031, Lots A00143/I and A00143/J), and dilutions into TAP or TP medium were made from a 10-mM stock in DMSO. Paromomycin (Sigma or EMD Millipore) and Zeocin (InvivoGen) were used at 10 μg/ml to select for and maintain strains that were transformed with constructs containing resistance markers.

Genetic crosses were performed essentially as described previously (60, 116, 117). When necessary, segregants were genotyped based on known phenotypes (LatB sensitivity, selectable marker, fluorescence, etc.) or by allele-specific PCR (60) using appropriate primers.

### Plasmids and transformation

pEB1-mNG (expressing EB1 protein fused to mNeonGreen) was a kind gift from Karl Lechtreck (68). Construction of pMO431 (*P*_*H/R*_:*MYO2-CrVenus-3FLAG*) was described previously (56); it expresses MYO2 tagged at its C-terminus with CrVenus-3FLAG from the hybrid *HSP70A*/*RBCS2* promoter (*P*_*H/R*_). All other plasmids used in this study were constructed using one-step isothermal assembly (118); synthetic DNA fragments and primers were obtained from Integrated DNA Technologies. All plasmids and corresponding sequence files are available through the Chlamydomonas Resource Center (https://www.chlamycollection.org). pMO654 (*P*_*H/R*_:*Lifeact-mNG*) was constructed by replacing *CrVenus*-*3FLAG* in pMO459 (57) with *mNG* from pEB1-mNG. A similar replacement of *CrVenus* in pMO611 (57) with *mNG* yielded pMO665 (*P*_*H*/*R*_*:mNG-3FLAG*), and genomic DNA sequences of the *PMH1* (Cre03.g164600), *MYO1* (Cre16.g658650), and *MYO3* (Cre13.g563800) coding regions (start codon to last coding codon, including all introns) were inserted into the *Hpa*I site of pMO665 to generate pMO683 (*P_H/R_:PMH1-mNG-3FLAG*), pMO668 (*P*_*H/R*_*:MYO1-mNG-3FLAG*), and pMO669 (*P*_*H*/*R*_*:MYO3-mNG-3FLAG*), respectively. Transformation by electroporation was done using a NEPA21 square-pulse electroporator and CHES buffer, as described previously (57), and transformants with strong Venus or mNG expression were identified by screening using a Tecan Infinite 200 PRO microplate reader at excitation and emission wavelengths of 515 and 550 nm, as described previously (57).

### Cell-cycle synchronization

Three different methods were used for cell-cycle synchronization in this study: (i) the 12L:12D/liquid TP method was essentially as described by Fang et al. (119) except that TP medium at 26°C was used in place of HSM; (ii) the 12L:12D/TAP agar method was as described previously (120), except that it was carried out at 26°C; and (iii) the -N method was exactly as described previously using a combination of 21°C and 33°C (121). Although overall synchrony and the timing of mitosis and cytokinesis as determined by microscopic examination varied slightly depending on the method, we observed no significant qualitative or quantitative difference in the cells’ response to F-actin perturbation introduced by LatB addition before the onset of cytokinesis.

### Light microscopy

Fluorescence and DIC microscopy of cells expressing Venus-, mNG-, and/or GFP-tagged proteins was performed as follows. Cells were mounted on a thin pad of TAP medium containing 1.5% low-melting-point agarose (Invitrogen) and sealed with a coverslip and VALAP. When desired, LatB was added to the agarose-containing medium at 3 μM. The cells were observed using a Nikon Eclipse 600-FN microscope equipped with an Apochromat x100/1.40 N.A. oil-immersion objective lens, an ORCA-2 cooled CCD camera (Hamamatsu Photonics), and Metamorph version 7.5 software (Molecular Devices). Signals of all fluorescent proteins were captured using YFP filters; chlorophyll autofluorescence was captured using Texas Red filters. For time-lapse experiments, the stage temperature was maintained at ~26°C using a heater (AmScope), and the slide was continuously illuminated by red LED lights to support photosynthesis. The distance from the leading edge of the cleavage furrow to the opposite side of the cell was determined at each time point to provide an approximate quantitative measure of the progress of furrow ingression. To visualize the rates of furrow ingression, local polynomial regression curves were generated using the LOESS (locally estimated scatterplot smoothing) method, and means and 95% confidence intervals of 1000 such curves were calculated using the loess.boot() function in the R package spatialEco. Images were post-processed using ImageJ (National Institutes of Health) and Photoshop (Adobe) software. Images from a single experiment with a single strain were processed identically and can be compared directly in the Figures.

The bright-field images in Fig. 4A and Fig. S2D were captured using a Leica DMI 6000 B microscope equipped with a x40 objective lens and Leica DFC 450 camera. Cell-wall removal by autolysin was performed essentially as described previously (122).

### Western blotting

Whole-cell extracts were prepared as described previously (61). SDS-PAGE was performed using Tris-glycine gels (8% for myosins, 11% for actin). After the proteins were transferred onto PVDF membranes, the blots were stained with a mouse monoclonal anti-FLAG (Sigma, F1804) or anti-actin (clone C4, EMD Millipore, MAB1501) antibody, followed by an HRP-conjugated anti-mouse-IgG secondary antibody (ICN Pharmaceuticals, 55564).

### Phylogenetic analysis

The amino-acid sequences of all myosins in the genomes of *Chlamydomonas* (MYO1, Cre16.g658650.t1.1; MYO2, Cre09.g416250.t1.1; MYO3, Cre13.g563800.t1.1 from Phytozome v5.5: phytozome.jgi.doe.gov)*, Arabidopsis thaliana* (40)*, D. melanogaster* (40), and *S. cerevisiae* (Myo1, YHR023W; Myo2, YOR326W; Myo3, YKL129C; Myo4, YAL029C; Myo5, YMR109W from Saccharomyces Genome Database: www.yeastgenome.org) were aligned using ClustalW 2.1 and the BLOSUM62 matrix. This alignment was then used to generate an unrooted Maximum Likelihood tree with approximate likelihood test using the VT model in PhyML (123).

## Acknowledgments

We thank Arthur Grossman, Frej Tulin, Kresti Pecani, Geng Sa, Howard Berg, Heather Cartwright, David Ehrhardt, Takako Kato-Minoura, Ritsu Kamiya, Martin Jonikas, Luke Mackinder, Karl Lechtreck, Prachee Avasthi, Alex Paredez, and Ryuichi Nishihama for valuable discussions, the provision of valuable reagents, or both. We also thank the Chlamydomonas Resource Center for providing essential strains and reagents. This work was supported by National Science Foundation Grants EAGER 1548533 and MCB 1818383 (to J.R.P.), and MCB 1515220 (to JGU), National Institutes of Health Grants R01GM126557 (to JGU) and 5R01GM078153 (to FRC).

---

**Fig. S1.**
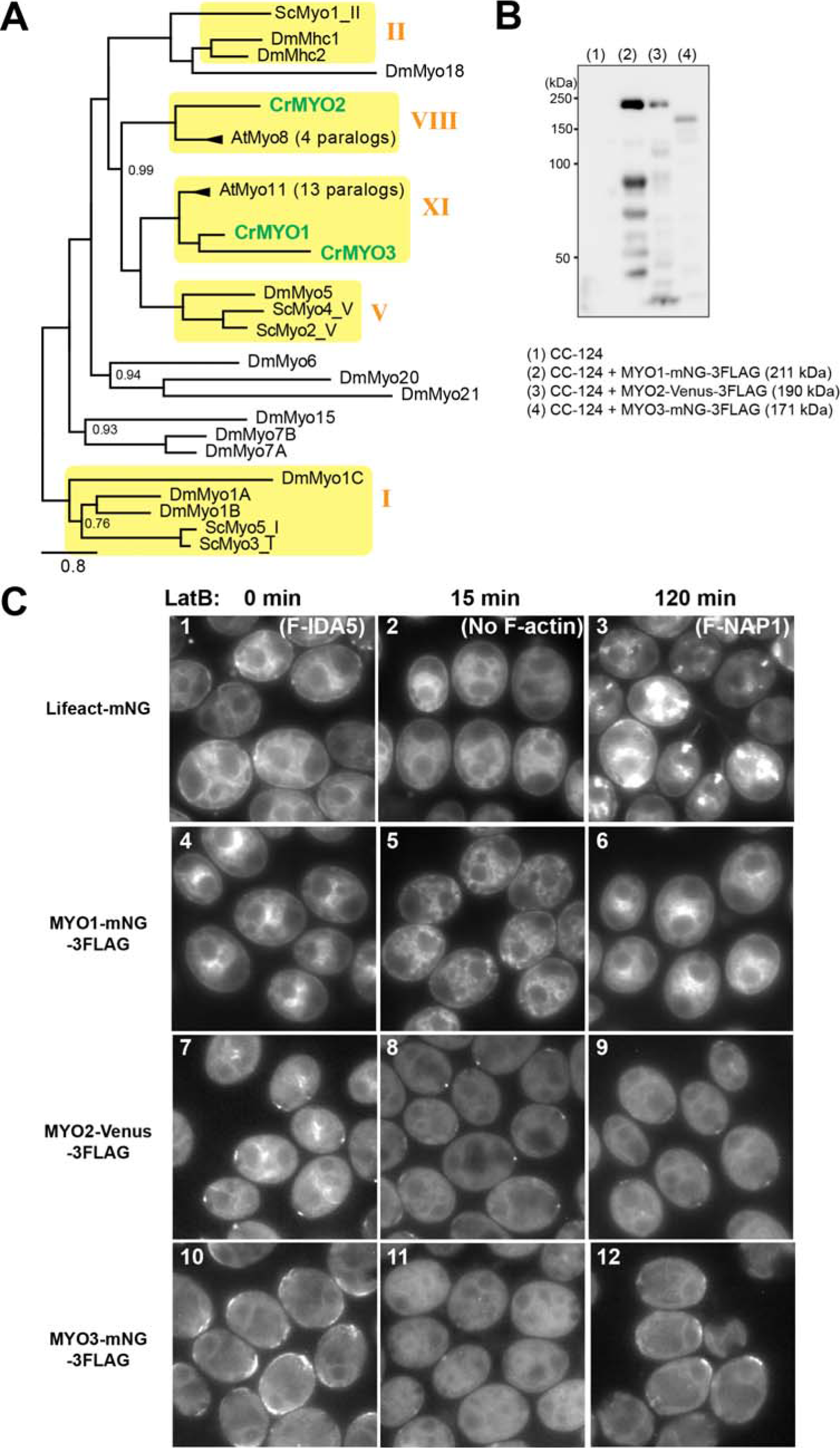
(*A*) Maximum-likelihood tree of amino-acid sequences of all myosins from *D. melanogaster* (Dm), *S. cerevisiae* (Sc), *A. thaliana* (At), and *C. reinhardtii* (Cr). Three *Chlamydomonas* myosins (green) clustered with plant-specific type-VIII and type-XI myosins. Scale bar, substitutions per residue. Branch supports below 1 are shown next to nodes. See Materials and Methods for details. (*B*) Expression of mNG-3FLAG-tagged or Venus-3-FLAG-tagged myosins. Whole-cell extracts from wild-type (CC-124) and transformed cells were analyzed by Western blotting using an anti-FLAG antibody. The positions of molecular-weight markers and the predicted molecular weights of the tagged proteins are indicated. (*C*) Response of F-actin and the tagged myosins to LatB treatment. Cells expressing Lifeact-mNG (1–3) or a tagged myosin (4–12) were treated with 10 μM LatB for the indicated times. See text for details.

**Fig. S2.**
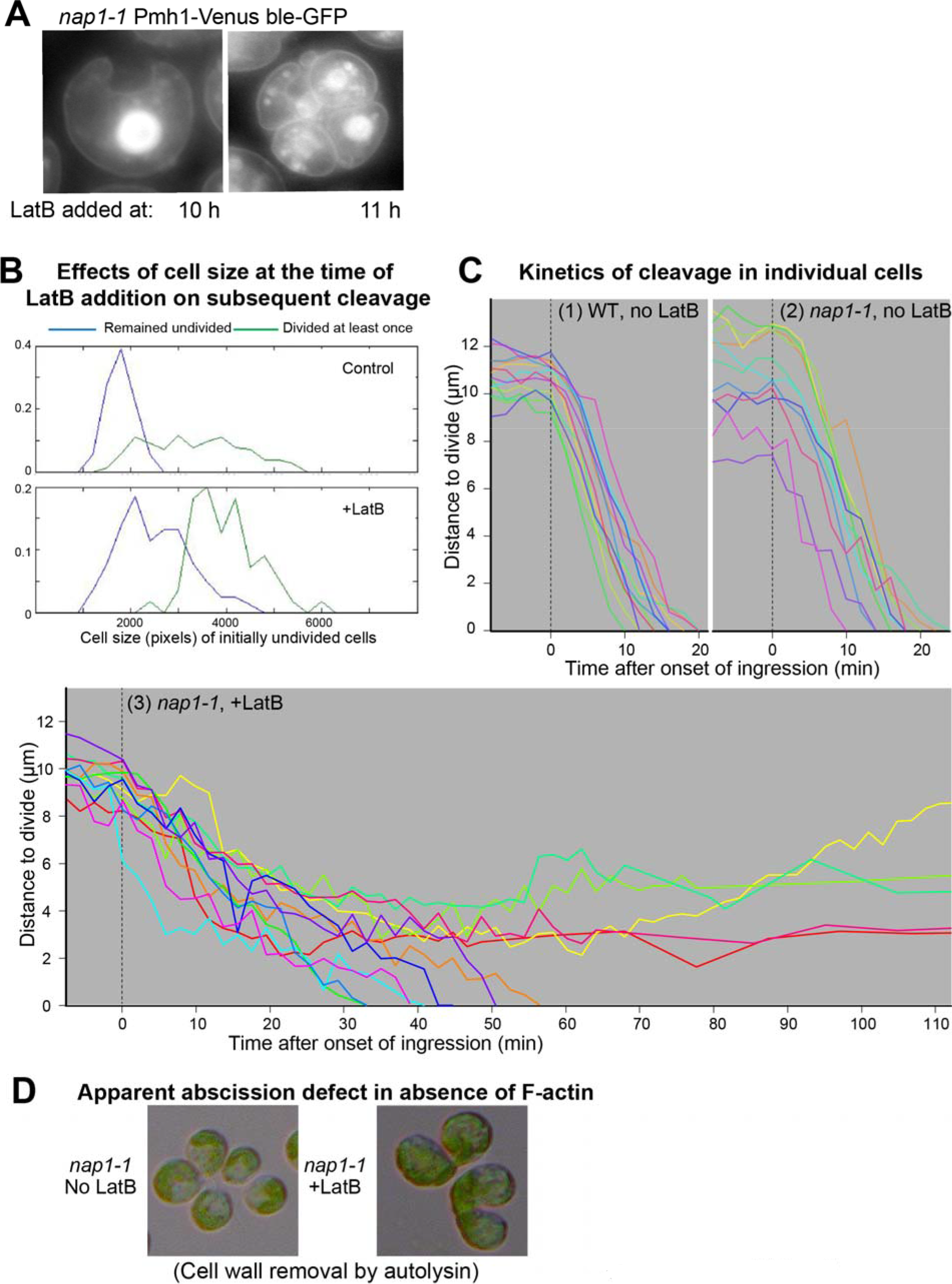
(*A*) Failure of mitosis when F-actin is eliminated sufficiently early in the cell cycle. In an experiment like that of Fig. 4*A*, *nap1-1* cells expressing PMH1-Venus and the nuclear marker ble-GFP were treated with 3 μM LatB beginning at the indicated times and imaged at 20 h. (*B*) Correlation between cell size and successful formation of furrows in the absence of F-actin. *nap1-1* cells were synchronized using the -N method (see Materials and Methods). At 11 hr, as cells began to enter divisions, they were transferred to fresh plates with or without 3 μM LatB and imaged every 30 minutes by brightfield time-lapse microscopy. The fractions of initially undivided cells that divided successfully were plotted as a function of the size of the cells at the time of transfer. (*C*) The kinetics of cleavage in individual cells in the experiments of Fig. 5. (*D*) Additional evidence for an abscission defect in cells lacking F-actin. In the experiment of Fig. 4*A*, control cells and cells treated with LatB beginning at 11 h were examined at 20 h. Cells were treated with autolysin before examination (see Materials and Methods). Control cells became rounder than normal as a result of cell-wall removal. ~5% of the LatB-treated cells remained connected by narrow intercellular bridges.

